# Spatial control of ARGONAUTE-mediated RNA silencing in anther development

**DOI:** 10.1101/2022.09.27.509800

**Authors:** Hinako Tamotsu, Koji Koizumi, Alejandro Villar Briones, Reina Komiya

## Abstract

Argonaute protein (AGO) in association with small RNAs is the core machinery of RNA silencing, an essential mechanism for precise development and defense against pathogens in many organisms. Here, we identified two AGOs in rice anthers, AGO1b and AGO1d, that interact with phased small interfering RNAs (phasiRNAs) derived from numerous long non-coding RNAs. Moreover, 3D-immunoimaging and mutant analysis demonstrated that rice AGO1b and AGO1d redundantly and cell type-specifically regulate anther development by acting as mobile carriers of these phasiRNAs from the somatic cell layers to the germ cells in anthers. Our study also highlights a new mode of reproductive RNA silencing via the specific nuclear and cytoplasmic localization of three AGOs, AGO1b, AGO1d, and MEL1, in rice pollen mother cells.

## Introduction

Site specificity of reproductive Argonaute proteins (AGOs) is essential for accurate germline development via transposable element silencing in many species ^1-4^. MEIOSIS ARRESTED AT LEPTOTENE1 (MEL1) is a germ cell-specific AGO among 19 AGO family members in rice ^5^. MEL1 interacts with 21-nucleotide (nt) phasiRNAs that have cytosine at the 5′-terminal position (C-phasiRNAs), which are derived from more than 1300 long non-coding RNAs named *21PHAS*s during pre-meiosis and early meiosis in rice ^6^. During plant reproduction, numerous 21- and 24-nt phasiRNAs are enriched in the anthers, the male organ in plants, via cleavage of miR2118/2275 and processing by Dicer-like proteins ^7-12^. *trans*-acting cleavage of 21-nt phasiRNAs in the cytoplasm of male germ cells has been detected ^13, 14^. In addition, 24-nt meiotic phasiRNAs and OsRDR6-dependent 24-nt small RNAs influence CHH DNA methylation during meiosis, leading to *cis* regulation by 24-nt phasiRNAs for maintaining normal meiosis ^15, 16^. Moreover, a recent study revealed that 24-nt siRNAs move from the tapetum to the meiocytes ^17, 18^, implying non-cell-autonomous regulation via small RNAs in anthers. Since somatic anther wall development affects germ cell development, their synchronization in anther development is indicative of the importance of cell-to-cell interaction ^19-21^. Intercellular communication via mobile signals is critical for cell fate determination in higher plants ^22, 23^. However, the non-cell-autonomous mechanism in anther development and the meiotic functions of 21-nt phasiRNAs remain unknown, although MEL1 interaction with 21-nt C-phasiRNAs plays an important role in the pairing between homologous chromosomes during early meiosis in rice ^6^. Therefore, focusing on reproductive AGOs as interactors of these phasiRNAs may illuminate the molecular roles of phasiRNAs in male organ development and in enhancing reproductive competence via the non-cell-autonomous developmental system in many species.

AGO1 generally binds to miRNA to execute post-transcriptional gene silencing in land plants ^24^. There are four members (AGO1a–d) in the AGO1 subfamily in rice, and AGO1a, b, and c bind to miRNAs at vegetative stages ^25^, as does *Arabidopsis* AtAGO1 ^26^. Rice *AGO1b/c/d* expression increases specifically during reproduction, while *AGO1a* is enriched at the vegetative stage. Our recent proteome and mRNA localization analyses have identified AGO1b and AGO1d as candidate miR2118-dependent soma AGOs ^27^, suggesting different functions of rice AGO1 subfamily proteins between vegetative and reproductive stages.

In this study, we identified phasiRNAs loaded onto rice AGO1b and AGO1d through small RNA-immunoprecipitation (RIP) and examined the role of AGO1b and AGO1d in anther development. We also successfully performed three-dimensional (3D)-immunostaining using whole anthers, which enabled us to distinguish the cell types and to identify the specific subcellular localization of AGO1b/d in each somatic cell layer and germ cell. Based on these results, we demonstrated the site-specific regulation of three AGOs, AGO1b/d and MEL1, in the anther phasiRNA pathway in rice.

## Results

### Seed sterility with abnormal anther development via *AGO1b* and *AGO1d* mutation

To elucidate the reproductive roles of AGO1d and AGO1b, we performed genome editing of *AGO1b* (Os04g0566500) or *AGO1d* (Os06g0729300) with the CRISPR/CAS9 system, using unique sequences of the 1^st^ exon as a guide RNA. We obtained three different allele mutants for each of *AGO1b* and *AGO1d*, namely *ago1b-1, ago1b-2, ago1b-3, ago1d-1, ago1d-2*, and *ago1d-3* (Fig. 1a; Supplementary Fig. 1). Each of the six mutants showed slightly reduced seed fertility compared to that in the *Japonica* rice ‘Nipponbare’ control, with no obvious effects in the mature anthers or pollen of any individual mutant (Supplementary Fig. 2). *ago1b-1* and *ago1d-3* were used for the following analysis. Using Wes as an automated western analysis system, we found a specific reduction of AGO1b protein in *ago1b-1* mutants, which have a 2-bp CC deletion in the 1^st^ exon of the *AGO1b* genomic region. In the *ago1d-3* mutants, with both a 20-bp deletion and a T substitution, AGO1d protein was specifically decreased, but not AGO1b (Fig. 1b). We next also generated double mutants of AGO1b/d in which *ago1b-1* pollen was used to pollinate the *ago1d-3* plant. Both AGO1b and AGO1d proteins were reduced in *ago1b-1 ago1d-3* double mutants (Fig. 1b). *ago1b-1* single mutant backcrossed once with Nipponbare showed almost normal fertility when compared to the fertility of segregating wild type (WT). In contrast, *ago1d-3* backcrossed once with Nipponbare showed partial sterility. Interestingly, *ago1b-1 ago1d-3* double mutants exhibited severe sterility compared to segregating WT (Fig. 1c), indicating the redundancy of AGO1b and AGO1d functions. Furthermore, the anthers of *ago1b-1 ago1d-3* double mutants were more curled and shorter, and with more varied sizes and shapes, than WT anthers (Fig. 1d, e). Mature pollen retained in the abnormal anthers lacked starch in severe-type anthers of this double mutant, while some pollen retained in the semi-abnormal anthers was stained (Fig. 1f, g; Supplementary Fig. 3). The mutant analysis demonstrates that AGO1b/d redundantly regulate anther development, thus affecting the maturation of pollen directly or indirectly.

**Figure 1.**
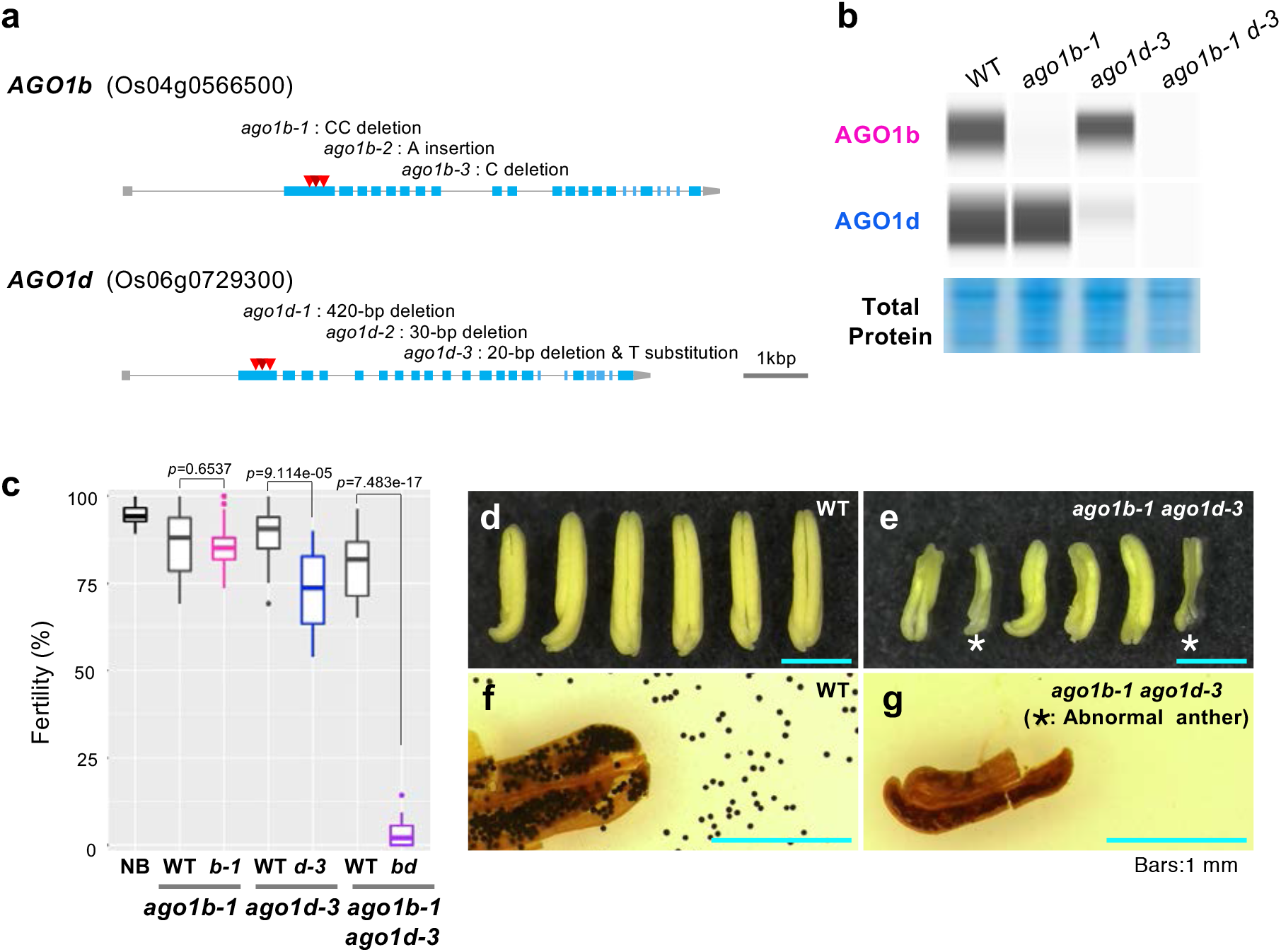
Seed sterility with abnormal anther development in *ago1b-1 ago1d-3* double mutants. **a**. Schematic structures of *AGO1b* and *AGO1d* genes, and three alleles for each of the *ago1b* and *ago1d* genome-editing mutants. Red triangles represent the deletion or insertion loci of *ago1b-1, ago1b-2, ago1b-3, ago1d-1, ago1d-2*, and *ago1d-3*. **b**. Wes analysis of AGO1b and AGO1d in Nipponbare (WT), *ago1b-1, ago1d-3*, and *ago1b-1 ago1d-3* double mutants. Total proteins were extracted from 0.5-mm anthers and are stained with CBB in the bottom panel. **c**. Fertility of Nipponbare (NB), WT segregated from *ago1b-1* backcrossed once with NB, *ago1b-1*, WT segregated from *ago1d-3* backcrossed once with NB, *ago1d-3*, WT segregated from *ago1d-3* backcrossed once with *ago1d-*, and *ago1b-1 ago1d-3* mutants. Data shown in box plots are from more than three biological replicates. Student’s *t* test. **d, e**. Mature anthers of WT and *ago1b-1 ago1d-3* double mutants. * indicates severely abnormal anthers with varied sizes and shapes. **f, g**. Mature pollen grains of WT and *ago1b-1 ago1d-3* double mutants stained with iodine-potassium iodide.

### Interaction of 21-nt U-phasiRNAs with AGO1b and AGO1d

Wes using anthers revealed that both AGO1b and AGO1d were enriched from the early meiosis stage to the post-meiosis stage among six stages of 0.4–0.9-mm anther development (Fig. 2a); each stage is explained in relation to germ and soma development in Supplementary Table 1. Next, to identify the small RNAs interacting with AGO1b and AGO1d, we performed RIP using 0.4–0.9-mm anthers from pre-meiosis to post-meiosis. AGO1b and AGO1d were identified using RIP fractions in Wes analysis, but they are difficult to detect in SDS-PAGE due to the low abundance of AGO1b/d complexes in anthers (Fig. 2b). Mass spectrometry analysis confirmed that AGO1b was immunopurified in AGO1b-RIP fractions, and AGO1d in AGO1d-RIP fractions, using antibodies generated in rabbits and mice, respectively (Supplementary Table 2). Next, the RNAs extracted from each of the AGO1b/d complexes were sequenced (Supplementary Table 3). AGO1b and AGO1d bound mainly to 21-nt small RNAs that have 5′-terminal uracil (U-phasiRNAs) (Fig. 2c, d). Additionally, the majority of AGO1b/d–small RNAs are categorized as 21-nt phasiRNAs derived from *21PHAS*s, which are anther-specific long non-coding RNAs (Fig. 2e). miRNAs interacting with AGO1b/d occurred at low frequency: 4.2% for AGO1b and 5.6% for AGO1d. In the AGO1b/d–miRNA group, AGO1d tended to bind to the 18 family members of miR2118, from miR2118a to miR2118r, more so than AGO1b (Fig. 2f). Argonaute proteins are known to sort small RNAs having specific 1^st^ nucleotides in plants ^26, 28^. The results of analysis of 21-nt U-phasiRNAs loading onto anther AGO1b/d coincide with the size and 5′-terminal nucleotide frequency of small RNAs interacting with AtAGO1; however, reproductive AGO1b/d–phasiRNAs are clearly distinct from the AtAGO1-miRNAs and vegetative OsAGO1 subfamily–miRNAs (Fig. 2) ^25, 26^.

**Figure 2.**
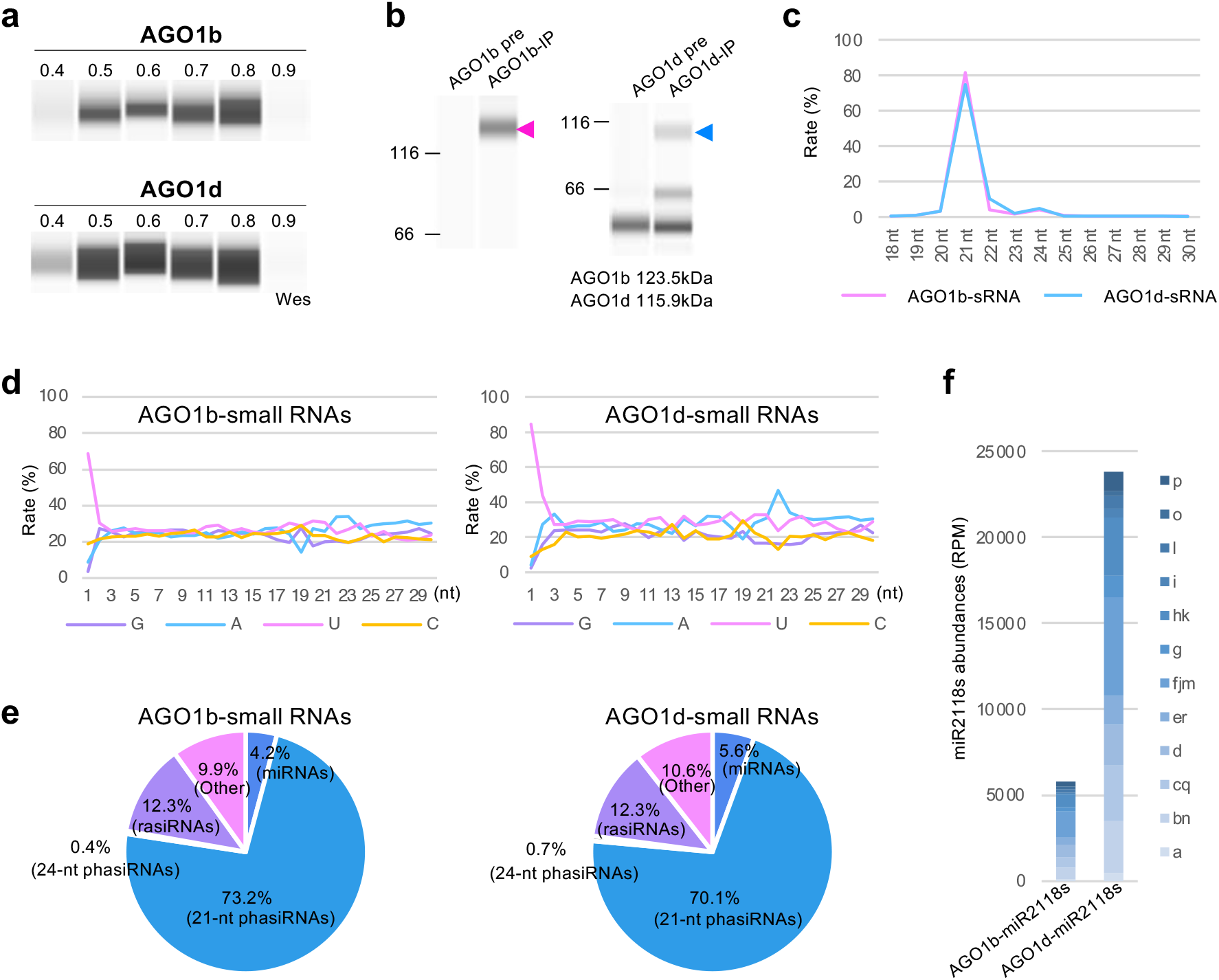
Association of 21-nt U-phasiRNAs with AGO1b and AGO1d. **a**. Wes analysis of AGO1b and AGO1d proteins in developing anthers. AGO1b and AGO1d total proteins, extracted from 0.4-mm (stage 1), 0.5-mm (stage 2), 0.6-mm (stage 3), 0.7-mm (stage 4), 0.8-mm (stage 5), and 0.9-mm (stage 6) anthers, were enriched during meiosis and post-meiosis (stages 2–5). Stage 1 is pre-meiosis, or primordial germ cell initiation; stages 2–4 are during meiosis; stage 5 is the microspore development stage; and stage 6 is the bicellular pollen stage. **b**. Wes images of AGO1b/d RIP fractions with anti-AGO1b or AGO1d. Arrowheads show the size distribution of AGO1b protein (magenta) and AGO1d protein (blue). **c**. Size distribution of AGO1b–small RNAs and AGO1d–small RNAs. **d**. Relative frequency of each nucleotide in the AGO1b/d–small RNAs. **e**. Pie charts summarizing the source of AGO1b/d–small RNAs. AGO1b and AGO1d mainly associate with 21-nt U-phasiRNAs derived from *21PHASs*/lncRNAs. **e**. Read counts of miR2118 family members in AGO1b–small RNAs and AGO1d–small RNAs. The reads data show the average counts of two replicates for AGO1b–small RNAs and four replicates for AGO1d–small RNAs.

### Segregation of U-phasiRNAs between AGO1b and AGO1d

Next, we performed clustering analysis using the *PHASIS* program ^29^ to detect the genomic loci of *21PHASs*, which are the origin of 21-nt phasiRNAs loaded onto AGO1b/d. Thus, we identified 2,337 clusters of AGO1b-phasiRNAs and 2,441 clusters of AGO1d-phasiRNAs in the rice genome (Fig. 3a; Supplementary Tables 4, 5). In contrast to the 21-nt phasiRNA clusters, we found 44 clusters in AGO1b–24-nt phasiRNAs and 73 clusters in AGO1d–24-nt phasiRNA (Fig. 3a; Supplementary Tables 6, 7). In the 21-nt phasiRNA clusters, the majority of AGO1b–phasiRNA clusters overlapped the loci of AGO1d-phasiRNA clusters (Fig. 3a, b). Interestingly, heatmap analysis enables us to classify the phasiRNAs into mainly two types of clusters, AGO1b-type and AGO1d-type clusters (Fig. 3c). AGO1b interacts predominantly with 21-nt phasiRNAs in the AGO1b-type clusters, while AGO1d interacts predominantly with those in the AGO1d-type clusters (Fig. 3c–e), suggesting that AGO1b/d also segregate subpopulations of the U-phasiRNAs.

**Figure 3.**
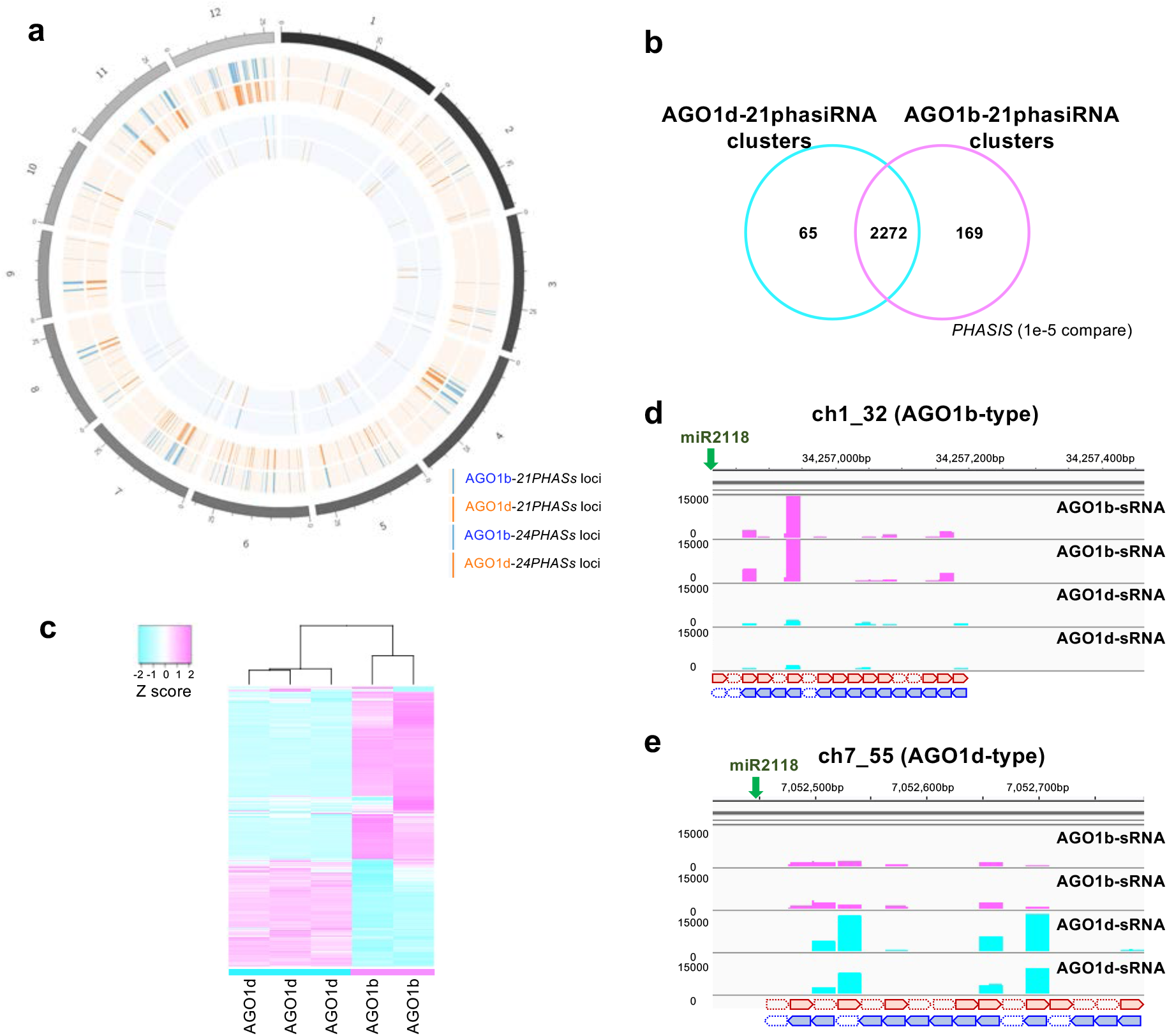
AGO1b and AGO1d segregation in U-phasiRNA sorting. **a**. Circular plot showing the distribution of the *21PHASs* or *24PHASs* on the 12 rice chromosomes from which 21-nt or 24-nt phasiRNAs interacting with AGO1b or AGO1d were derived. *21PHASs* loci, where 21-nt phasiRNAs associating with AGO1b (blue) and AGO1d (orange) are produced, are displayed in the outer pale orange zone. *24PHASs* loci, where 24-nt phasiRNAs associating with AGO1b (blue) and AGO1d (orange) are produced, are displayed in the inner pale blue zone. **b**. Venn diagram of overlapping 21-nt phasiRNA clusters, *21PHASs* genomic loci that were identified using small RNAs that sorted into AGO1b or AGO1d. **c**. Heatmap of predominant pattern of small RNAs interacting with AGO1b and AGO1d, and total small RNAs extracted from 0.5-mm anthers. **d, e**. Small RNA-seq reads of AGO1b–small RNAs and AGO1d–small RNAs (two replicates of each) that were mapped to the AGO1b-type clusters (**d**) and AGO1d-type clusters (**e**).

### Nuclear AGO1b and cytoplasmic AGO1d in soma and germ

The anther is a major part of the male reproductive organ in plants, consisting of somatic anther walls and germ cells, known as the pollen. The somatic anther walls consist of four cell layers (epidermis, endothecium, middle layer, and tapetum) during early meiosis, in which stages AGO1b and AGO1d are abundant (Fig. 2a; Supplementary Table1). Recently, we developed the 3D-anther immunostaining system at subcellular and single-cell resolution ^30^. To investigate the localization of these AGO1b/d proteins in anthers, we performed the 3D multiple immunostaining against AGO1b and AGO1d using whole-mount anthers at early meiosis. We captured continuous image data of the anther, consisting of 80 slices with 0.6-μm intervals from the outer epidermis to the inner pollen mother cells in longitudinal Z sections, and created a movie by stacking these images (Movie 1). We thus were able to detect the localization of AGO1b/d proteins in each cell layer and in pollen mother cells of rice anthers at this stage (Movie 1; Fig. 4). It is also possible to distinguish the four somatic cell layers in X sections, which are cross-sections of anthers (Movie 1, upper figure).

**Figure 4.**
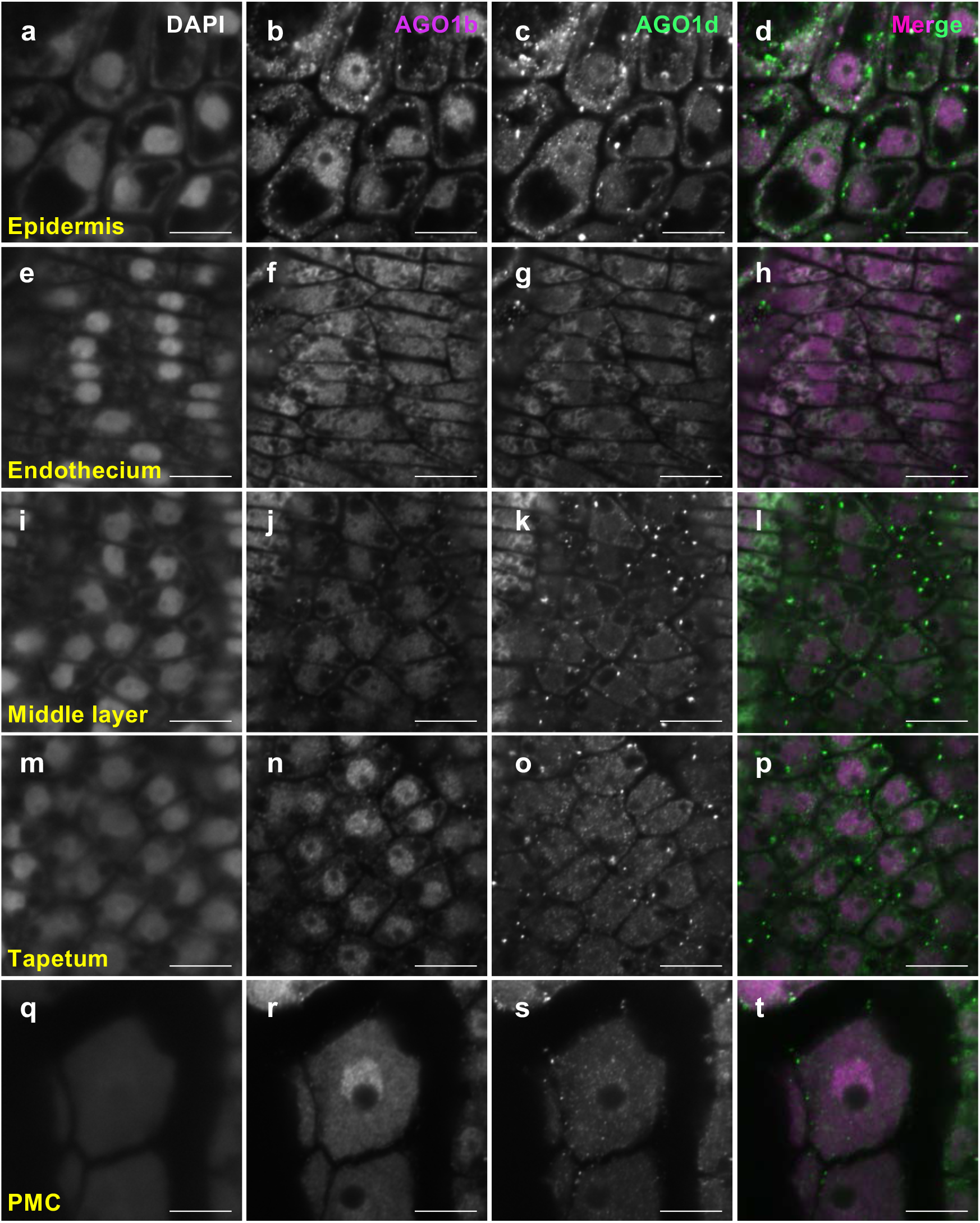
AGO1b nuclear localization in epidermis, tapetum, and pollen mother cells. **a–t**. Immunostaining using whole 0.5-mm anthers at early meiosis (stage 2) against AGO1b (magenta, **bfjnr**) and AGO1d (green, **cgkos)**. DAPI staining was used as a nucleolus marker (**aeimq**). AGO1b and AGO1d fluorescence images were merged (**dhlpt**). This (3D)-immunostaining enables us to distinguish the cell types as epidermis (**a–d**), endothecium (**e–h**), middle layer (**i–l**), tapetum layer (**m–p**), and pollen mother cells (PMC) (**q–t**). Laser excitation/emission are 405 nm/410–455 nm for DAPI, 488nm/490-552nm for AGO1d, and 561 nm/544–615 nm for AGO1b. Scale bars, 10 μm.

Both AGO1b and AGO1d were enriched in the nucleus and at the cell membrane of the epidermis, whose cells are the largest in the anther wall layers (Fig. 4a–d). In the elongated-transverse endothecium cells, AGO1d was more abundant in cytoplasm near the nucleus (Fig. 4e–h). In the cells of the middle layer, AGO1b was present in the nucleus. However, weak fluorescence signals of AGO1d were detected in the cytoplasm in the middle layer cells (Fig. 4i–l). Furthermore, AGO1b localization in the tapetum layer was restricted to the nucleus, while AGO1d was present in both nucleus and cytoplasm (Fig. 4m–p). In somatic anther walls, AGO1b thus tends to localize in the nucleus, and AGO1d in the cytoplasm. The differing intracellular localization of AGO1b in the nucleus and AGO1d in the cytoplasm of all four cell types in the anther wall concurs with the segregation of phasiRNA loading onto AGO1b and AGO1d (Figs. 3c–e, 4a–p; Movie 1). Furthermore, AGO1b was mainly detected in the nucleus of pollen mother cells at early meiosis (Figs. 4r, 5a), and AGO1d was present in both nucleus and cytoplasm in germ cells (Fig. 4s). In contrast, MEL1 is enriched in the cytoplasm of pollen mother cells (Fig. 5b, c) ^6^, suggesting that proper intracellular localization, in addition to cell specificity, among AGO1b/AGO1d/MEL1 is required for the germ cell development.

**Figure 5.**
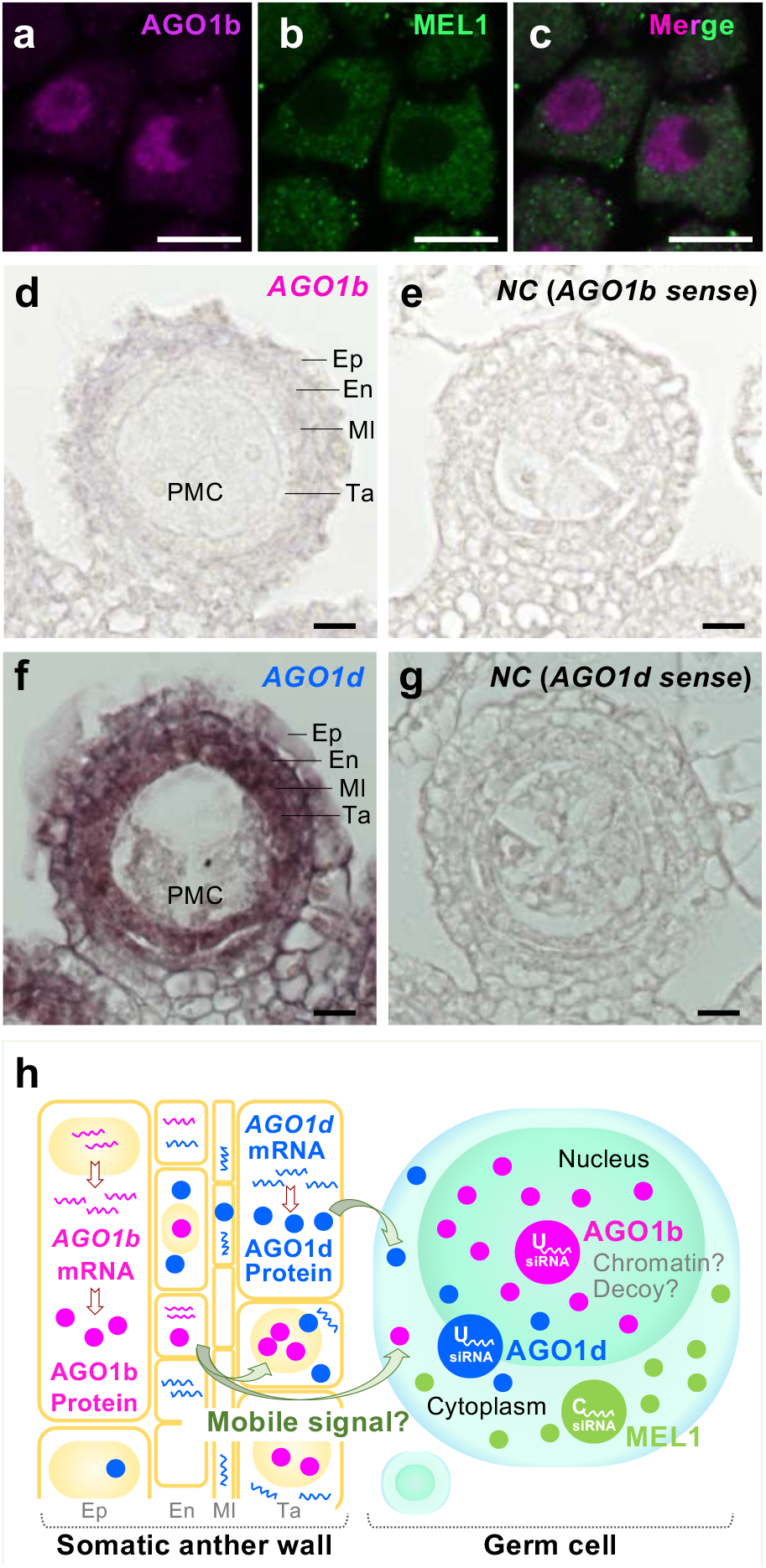
U-phasiRNAs-carrying AGO1b/d as mobile signals from outer anther wall cells to inner layers and pollen mother cells. **a–c**. Immunostaining using anti-AGO1b and anti-MEL1 in pollen mother cells of the 0.5-mm anther (early meiosis). AGO1b is enriched in the nucleolus (magenta) **(a)**, while MEL1 localization is restricted to the cytoplasm (green) **(b)**. Scale bars, 10 μm. **d–g**. *In situ* hybridization of AGO1b, AGO1d, and negative control (NC, sense strand) probes using anthers from 2.0–2.5-mm inflorescences, at stages 1 and 2. *AGO1b* localization was more highly enriched in epidermis (Ep) and endothecium (En) than tapetum layers (Ta) **(d)**. *AGO1d* expression is abundant in anther wall, especially tapetum (Ta), middle layer (Ml), and endothecium (En) **(f)**. Scale bars, 10 μm. **h**. Model of rice site-specific RNA silencing in rice anther development. *AGO1b* and *AGO1d* mRNA are increased in the somatic anther wall during pre-meiosis and early meiosis and translated; AGO1b/d proteins interacting with 21-nt U-phasiRNAs then move and become enriched in the tapetum and pollen mother cells, acting as a mobile signal. AGO1b/d–U-phasiRNAs are detected in both soma and germ, and thus differ from MEL1–C-phasiRNAs in germ. Moreover, their intracellular localization differs in pollen mother cells: cytoplasmic for MEL1, nucleolar for AGO1b, and both cytoplasmic and nuclear for AGO1d.

### U-phasiRNAs-carrying AGO1b/d as mobile signals from outer anther wall cells to inner layers and pollen mother cells

Next, we performed the *in situ* hybridization to investigate the RNA localization of *AGO1b* and *AGO1d* in anthers. *AGO1b/d* mRNAs are both detected in the epidermis, endothecium, and middle layers of somatic anther walls. *AGO1d* mRNA also occurs in the somatic anther walls, more markedly in the tapetum, during early meiosis (Fig. 5d– g). Moreover, *AGO1b/d* mRNAs are rarely detected in pollen mother cells (Fig. 5d–g), while *MEL1* is specifically expressed in germ cells ^*5*^.

The 3D-immunoimaging results also indicate that AGO1b is enriched in pollen mother cells in addition to being localized in the tapetum cell layer (Figs. 4nr, 5a). However, *in situ* hybridization showed that *AGO1b* mRNA was expressed in the outer layers of the anther wall, and not in the inner tapetum layers or pollen mother cells (Figs. 5d). Moreover, AGO1d was localized in the pollen mother cells, while *AGO1d* mRNA was strongly expressed in the tapetum cell layers (Figs. 4s and 5f). The different localization of mRNA and protein of AGO1b/d subfamily genes in anther suggests that AGO1b and AGO1d migrate from outer layers to inner layers and pollen mother cells as carriers of phasiRNA.

## Discussion

In summary, we have identified novel RNA silencing via AGO1b/d–U-phasiRNA complexes, which are enriched in both soma and germ in anthers, and specifically localized to the nucleus in pollen mother cells; this silencing differs from the germ-specific RNA silencing with cytoplasmic localization by MEL1–C-phasiRNAs (Fig. 5h). Of particular interest is the discrimination of subcellular localization in anther wall layers, where AGO1b is enriched in the nucleus and AGO1d is mainly localized in the cytoplasm. In addition to this segregation of AGO1b/d localization in soma, the differences between MEL1, AGO1b, and AGO1d in germ subcellular localization may be critical for the function of C-phasiRNAs and U-phasiRNAs. Recent studies, including the *trans*-acting silencing of MEL1–C-phasiRNAs via cleavage in germ ^13 14^, reflect the possibility that AGO1b/d–U-phasiRNAs in the nucleus act as decoys, similar to the function of *Arabidopsis* AGO10 ^31^. Alternatively, AGO1b/d–U-phasiRNAs may have a novel *cis*-epigenetic function during meiosis, as is the case for *cis*-regulation of 24-nt meiosis phasiRNAs with CHH DNA methylation ^16^. Our results on cell type- and intracellular-specific AGO localization in the rice male organ illuminate the importance of AGO combination-mediated RNA silencing during reproduction. Epigenetic regulation of anther-specific AGO2 precisely initiates the programmed cell death in tapetum developments, resulting in normal pollen development in rice ^32^. Furthermore, the phosphorylation or ubiquitination of reproductive AGOs are vital for temperature-dependent male fertility or meiosis to proceed the accurate reproduction via the phasiRNAs control in crops ^33 34^. Therefore, understanding reproductive AGOs functions and combination could lead to the solution of the phasiRNAs molecular roles in male organ development, which has critical implications for a stable yield of crops.

We found that the AGO1b/d proteins were abundant in tapetum and pollen mother cells, while *AGO1b* mRNA was lacking in these cell types and *AGO1d* mRNA was also absent in the pollen mother cells (Figs. 4, 5). *ago1b-1 ago1d-3* double mutants additionally exhibited defects of pollen accompanied by abnormal anther development (Fig. 1). Recent reproductive studies, the cell-to-cell transportation of 24-nt phasiRNAs and pollen-development regulation by the somatic transcription factors, imply the non-cell-autonomous regulation in anther development ^18, 35^. We therefore speculate that AGO1b/d constitute a mobile signal from the outer anther wall to the inner wall, tapetum, and pollen mother cells as carriers of U-phasiRNAs (Fig. 5h). Mobile protein signals including florigen and homeobox transcription factors are essential for accurate development and cell fate determination in higher plants ^36-38^. Understanding the nature of reproductive AGO–phasiRNA silencing via cell-to-cell communication should provide new insights into the non-cell-autonomous mechanism in anther development, and could greatly enhance the reproductive competence of plants.

## Materials and Methods

### Plant materials and growth conditions

Rice (*Oryza sativa L*., subspecies *Japonica*, cultivar Nipponbare) was used in this study. Plants were grown in growth chambers at 70% humidity under long day (LD) conditions, with a daily cycle of 14 h of light at 29.5–30 °C and 10 h of dark at 25 °C, for the mutant analysis in Fig. 1. To align the sampling stages, plants were grown for 40 days under LD conditions and then transferred to short day conditions (10 h of light and 14 h of dark) until harvest for the data in Figs. 2–5.

### Generation of AGO1b and AGO1d antibodies

A synthetic peptide, AGO1b (Cys-GSSQRAERGPQQH-OH), was used to raise rabbit polyclonal antibody against rice AGO1b. Oligopeptides AGO1d-1 (Cys-GRGSYYPQAQQYH-OH) and AGO1d-2 (Cys-HQQPYNSSVRPQH-OH) were used to raise antibodies in rabbits and mice. The rabbit antisera were purified by affinity chromatography.

### Western analysis

Wes is automated western analysis using capillary, not gel blotting. Results are shown in a digital image such as a western blotting analysis. Total proteins were extracted from 0.4–0.9-mm anthers of the WT (Fig. 1a), and 0.5-mm anthers of the WT and *ago1b-1* and *ago1d-3* mutants (Fig. 4b). The anthers were ground and mixed with extraction buffer (150 mM NaCl, 50 mM Tris–HCl (pH 7.5), 0.1% (v/v) Tween 20, 10% (v/v) glycerol, 1 mM dithiothreitol (DTT), 1 mM Pefabloc SC (Roche), 1× Complete Protease Inhibitor Cocktail (Roche)). After centrifuging twice to remove debris, total proteins were extracted. Anti-AGO1b/d (1/20 dilution) were used as primary antibodies.

### Small RNA-immunoprecipitation (RIP)

RIP was performed as previously described ^39^. Anthers (0.4–0.8 mm) were homogenized in extraction buffer (see above). AGO1b/AGO1d–small RNA complexes were washed four times in wash buffer (20 mM Tris–HCl (pH 7.5), 150 mM NaCl, 5 mM MgCl_2_, 5 mM DTT, 0.1% (v/v) NP-40, 1× Complete Protease Inhibitor Cocktail).

### Small RNA sequencing

NEXTflex small RNA-seq libraries (150 bp paired-end) were prepared (Illumina, Compatible) and sequenced using Illumina Novaseq. Four or two biological replicates were prepared for sequences of small RNAs binding to AGO1b or AGO1d (Fig. 2).

### Small RNA sequencing data analysis

Sequencing reads were trimmed using TrimGalore (http://www.bioinformatics.babraham.ac.uk/projects/trimgalore) to remove adapter sequences and sequencing bias with the following parameters: stringency = 5, quality = 20, and three prime clip R1 = 60. After trimming, reads that ranged from 18 bp to 30 bp were used. To annotate the small RNAs, perfectly matched reads were mapped onto the rice genome IRGSP1.0 (https://rapdb.dna.affrc.go.jp) with Bowtie, and the datasets of miRNA from miRbase (http://www.mirbase.org/ftp.shtml) and repeat data from RAP-DB (https://rapdb.dna.affrc.go.jp/) were used.

To identify 21- and 24-nt phasiRNA clusters, 18- to 30-nt trimmed small RNAs were analyzed using *PHASIS* ^29^ with default parameters. Significant clusters (*p*-value ≤ 1e^-5^) from the replicates were merged. Overlapping clusters among the samples were extracted using the phasmerge function in *PHASIS*.

### *In situ* hybridization (ISH)

ISH was performed as previously described ^40^. The primer sets for the probes are listed in Supplementary Table 8.

### Immunostaining using whole mounts of anthers

3D-immuno imaging was performed as described ^30^. Anthers fixed in 4% paraformaldehyde were transferred to PME buffer (50 mM PIPES, 5 mM EGTA and 5 mM MgSO_4_, pH 6.9) on a MAS-coated microscope slide, cut in distilled water using a scalpel, and incubated for 30 min at 25 °C. The cleaved anthers were then blocked with 3% bovine serum albumin (BSA) in PME for 60 min. The samples were placed in primary antibody solutions (rabbit anti-AGO1b or mouse anti-AGO1d, diluted 1/500 with 3% BSA in PME), degassed five times at 0.05 MPa for 2 min, and incubated overnight at 4 °C. After washing three times for 5 min with PME, the slide was placed in secondary antibody solution (Alexa Fluor 568-conjugated anti-rabbit IgG (Invitrogen, A11036) or Alexa Fluor 488-conjugated anti-mouse IgG (Invitrogen, A11001), diluted 1/200 with 3% BSA in PME), and degassed as above. The slide was incubated in a dark chamber for 2 h at room temperature, and then incubated overnight at 4 °C. The samples were washed three times with PME buffer for 5 min, incubated for 15 min at 25 °C in DAPI (Sigma, MBD0015), and washed again with PME buffer as above. Samples were mounted in ProLong Gold antifade reagent (Invitrogen, P10144). Images were captured using an LSM780 microscope (Carl Zeiss).

### Visualization of the 3D immunostaining of the entire anthers

The images was captured using an LSM780 microscope (Carl Zeiss). Conditions: 40x (1.4 oil) Plan Apochromat lens (Movie1) and 63x (1.46 oil) Plan Apochromat lens (Figs. 4, 5, Movie 1) for detection, 405, 488, and 561 nm laser lines for DAPI, Alexa Fluor 488, and Alexa Fluor 568 excitation, 410–455 nm (DAPI), 490–552 nm (Alexa Fluor 488) and 544–615 nm (Alexa Fluor 568) filter emission. The images and animation were created using the ZEN (Carl Zeiss) or Imaris 9 (Bitplane AG) software (Figs. 4, 5, Movie 1).

### Mass spectrometry

The protein samples, which were AGO1b/AGO1d-IP fractions, were reduced with DTT and then alkylated with iodoacetamide and digested overnight using Lys-C/Trypsin (1:50, enzyme to protein; Promega). After terminating the digestion with 1% trifluoroacetic acid, the peptide mixture was cleaned with desalting C18 tips (StageTip, Thermo Fisher Scientific), and subsequently vacuum-dried and dissolved in 0.1% Formic acid, 0.5% Acetic acid in water for LC/MS analysis.

Data were collected using an Orbitrap-Fusion Lumos mass spectrometer (Thermo Fisher Scientific) coupled with the Waters nanoACQUITY liquid chromatography system. A trap column (nanoACQUITY UPLC 2G-V/M Trap 5 µm Symmetry C18, 180 µm × 20 mm, Waters) and an analytical column (nanoACQUITY UPLC HSS T3 1.8 um, 75 µm × 150 mm, Waters) were used for chromatographic separation of samples. Peptides were separated at a flow rate of 500 nL/min using a gradient of 1–32% acetonitrile (0.1% formic acid) over 60 min. The CHOPIN method ^41^ was used along with the Orbitrap-Fusion mass spectrometer using Xcalibur (v.3.0; Thermo Fisher Scientific).

Raw data files were searched against a composite target/decoy database using SEQUEST (Proteome Discoverer, v.2.2, Thermo Fisher Scientific).

### Gene targeting construction and transformation

pZH_gYSA_MMCas9 and pU6gRNA-oligo were used as vectors ^42^. For editing of the *AGO1b* locus, 5′-AGCCATACTATGGCGGACCT-AGG (PAM)-3′ was used for the target sequence of the CRISPR/Cas9 vector as a single guide RNA. 5′-CTCCGGAGGCATCATCACCA-CGG (PAM)-3′ was used for genome editing of the *AGO1d* locus, and *ago1d-1* and *ago1d-2* were generated; 5′-GACCGTGGTGATGATGCCTC-CGG (PAM)-3′ was used for genome editing of the *AGO1d* locus, and *ago1d-3* was generated. Underlines show the PAM sequences. First, the oligos for guide RNAs were cloned into a pU6gRNA-oligo vector. Second, OsU6 promoter::gRNA was cut with *AscI* and *PacI*, and cloned into the pZH_gYSA_MMCas9 binary vector with hygromycin phosphotransferase selection ^42^. Transgenic rice plants were generated by *Agrobacterium*-mediated transformation of rice calli (cv. Nipponbare) ^43^.

## Data Availability

*AGO1b*; Os04g0566500, *AGO1d*; Os06g0729300; *MEL1*; Os03g0800200. The gene datasets used during the current study are available in the RAP-DB repository (https://rapdb.dna.affrc.go.jp/). RIP data have been deposited in the DNA Bank of Japan (DDBJ), under the accession code PSUB017586.

## Acknowledgments

This work was supported by the JST FORESTO Program (Grant Number JPMJFR204U, Japan), the JST PRESTO Program (Grant Number JPMJPR17Q3, Japan), the Naito Foundation, and the Okinawa Institute of Science and Technology Graduate University, Japan. We thank all members of the Science and Technology Group and FORESTO members (Siomi panel) for their helpful discussions.

## Author Contributions

R.K. conceived the study, conducted most of the data analysis, and wrote the manuscript. R.K. performed RIP and bioinformatics analysis of small RNAs. H.T. generated *ago1b, ago1d* and *ago1b ago1d* mutants, and performed mutant phenotyping. R.K. performed 3D-immunostaining, and K.K performed *in situ* hybridization. H.T. performed protein experiments. A.V.B. performed proteome analysis. H.T. and K.K. assisted in creating imaging figures and discussion.

## Competing Interests

The authors declare no competing interests.

## Additional information

Supplementary information tables and figures are available for this paper.

## Material and correspondence

Correspondence and material requests to Reina Komiya.

## Figure and Movie legends

**Movie 1.**
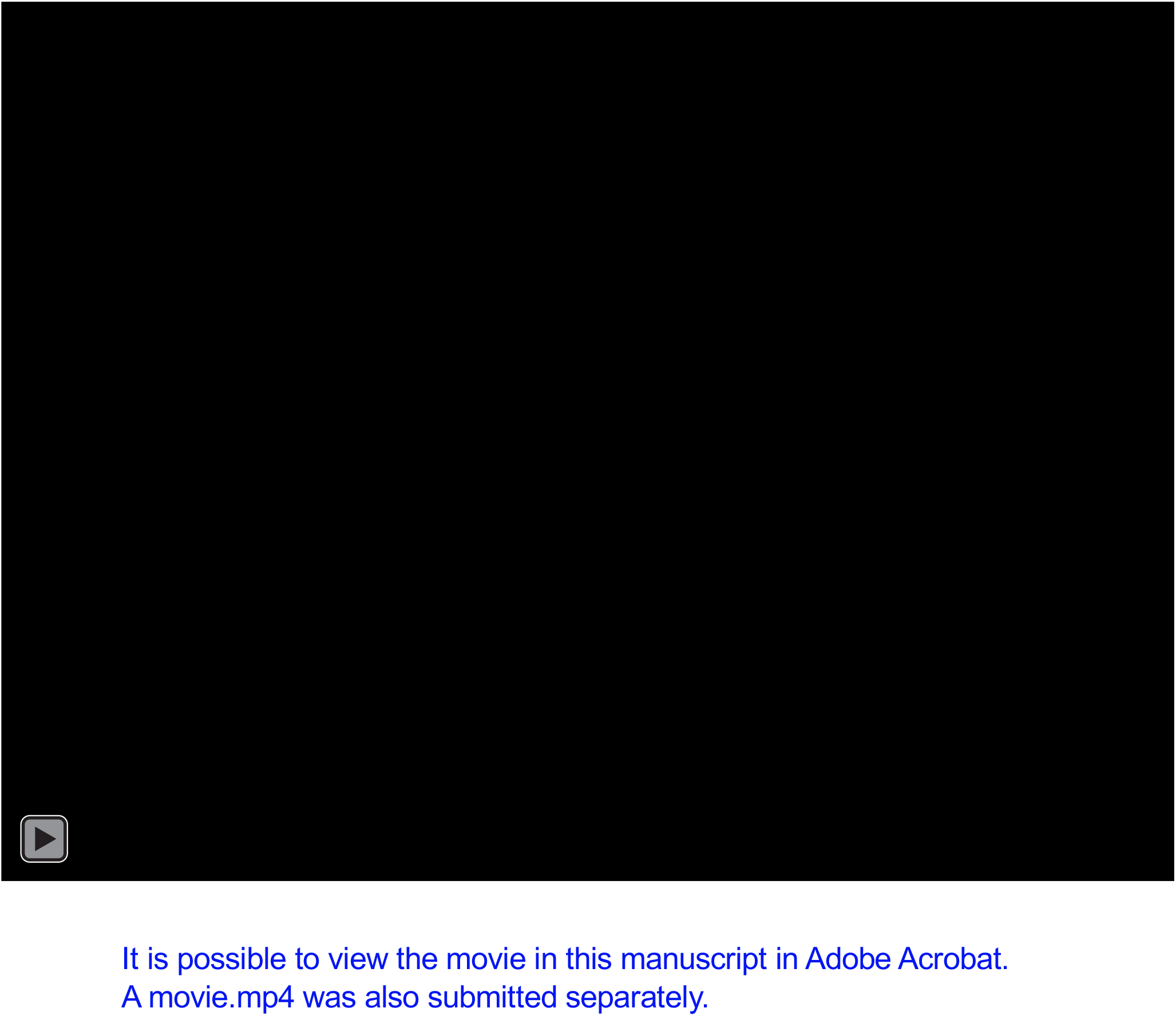
3D movie of AGO1b/d immunostaining using whole anthers. Cyan signals indicate DAPI staining. Magenta signals indicate the indirect fluorescence of the AGO1b protein. Green signals indicate the indirect fluorescence of the AGO1d protein. The 0.5 mm-long anthers at early meiosis were used for 3D-multiple immunoimaging. An upper right figure shows a cross-section (X section) of the 3D anther immunoimaging, in which 0.5-mm anthers from the early meiosis stage were used. Laser excitation/emission are 405 nm/410–455 nm for DAPI, 488nm/490-552nm for AGO1d, and 561 nm/544–615 nm for AGO1b. Scale bar, 10 μm.

**Supplementary Figure 1. a**. Schematic structure and alignment of the *AGO1b* gene and its mutants. We generated three mutant *ago1b* alleles. Three red triangles represent the deletion or insertion loci of *ago1b-1, ago1b-2*, and *ago1b-3*. Red boxes show deletions, and the blue box shows an insertion. **b**. Schematic structure and alignment of the *AGO1d* gene and two of its mutants. We obtained three mutant *ago1d* alleles. Three red triangles represent the deletion or substitution loci of *ago1d-1, ago1d-2*, and *ago1d-3*. The green box shows the sequence of a guide RNA.

**Supplementary Figure 2. a**. Fertility of Nipponbare (NB), *ago1b-1* (*b-1*), *ago1b-2* (*b-2*) *ago1b-3* (*b-3*), *ago1d-1* (*d-1*), *ago1d-2* (*d-2*), and *ago1d-3* (*d-3*). Data shown in box plots are from more than three biological replicates. Student’s *t* test. **b**. Mature anthers (left), pistil (middle), and pollen grains (right) of Nipponbare (control), *ago1b-1*, and *ago1d-3*. The single mutants, *ago1b-1* and *ago1d-3*, showed normal anthers and pistils compared to the those of Nipponbare. Furthermore, mature pollen grains of *ago1b-1* and *ago1d-3* were also stained with iodine-potassium iodide.

**Supplementary Figure 3. a**. Anthers of WT. **b**. Starch staining of pollen from WT anthers. **c**. Anthers of *ago1b-1 ago1d-3* double mutants with several abnormal shapes from severe to mild types (#1–6). **d–i**. Starch staining of pollen from a severely abnormal and semi-abnormal anthers of the *ago1b-1 ago1d-3* double mutant (#1–6). Non-staining pollens reflect the abnormality of pollen activity and development (**d and h**). Semi-abnormal anthers of the double mutant contain non-staining pollens (white arrows) as well as stained pollen grains (magenta arrows) (**e, f, g, and i**). Most of the stained pollen grains were trapped in the anthers of the *ago1b-1 ago1d-3* double mutant (**e, f, g, and i**), perhaps due to defects of somatic anther wall developments. Scale bars are 1 mm.

## Notes

### Competing Interest Statement

The authors have declared no competing interest.

