## Supplementary Figures for "Spatial control of ARGONAUTE-mediated RNA silencing in anther development"

Supplementary Figure 1

**a**

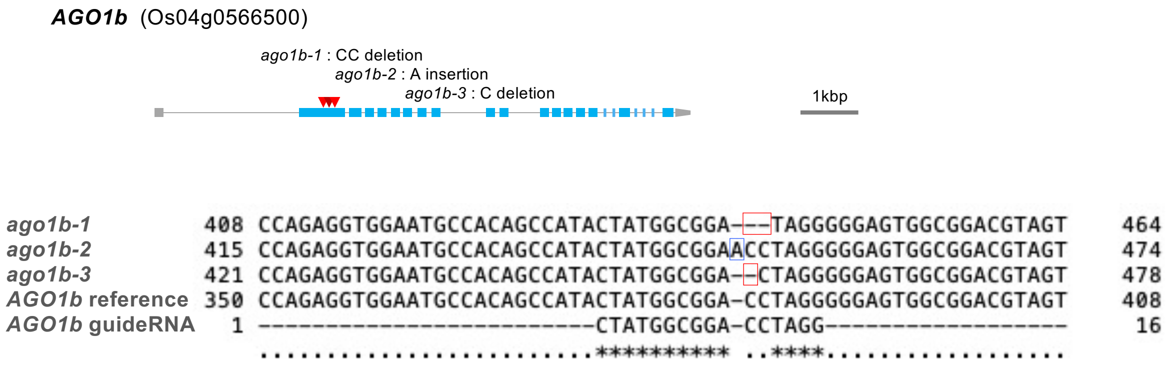


**b**

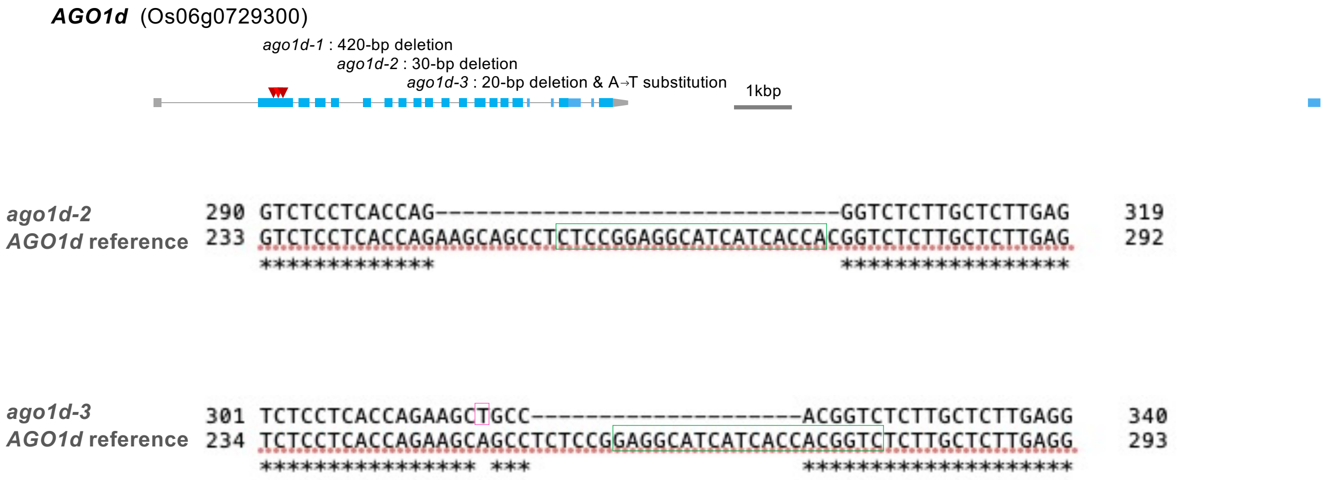


**Supplementary Figure 1. a**. Schematic structure and alignment of the *AGO1b* gene and its mutants. We generated three mutant *ago1b* alleles. Three red triangles represent the deletion or insertion loci of *ago1b-1*, *ago1b-2*, and *ago1b-3*. Red boxes show deletions, and the blue box shows an insertion. **b**. Schematic structure and alignment of the *AGO1d* gene and two of its mutants. We obtained three mutant *ago1d* alleles. Three red triangles represent the deletion or substitution loci of *ago1d-1*, *ago1d-2*, and *ago1d-3*. The green box shows the sequence of a guide RNA.

Supplementary Figure 2

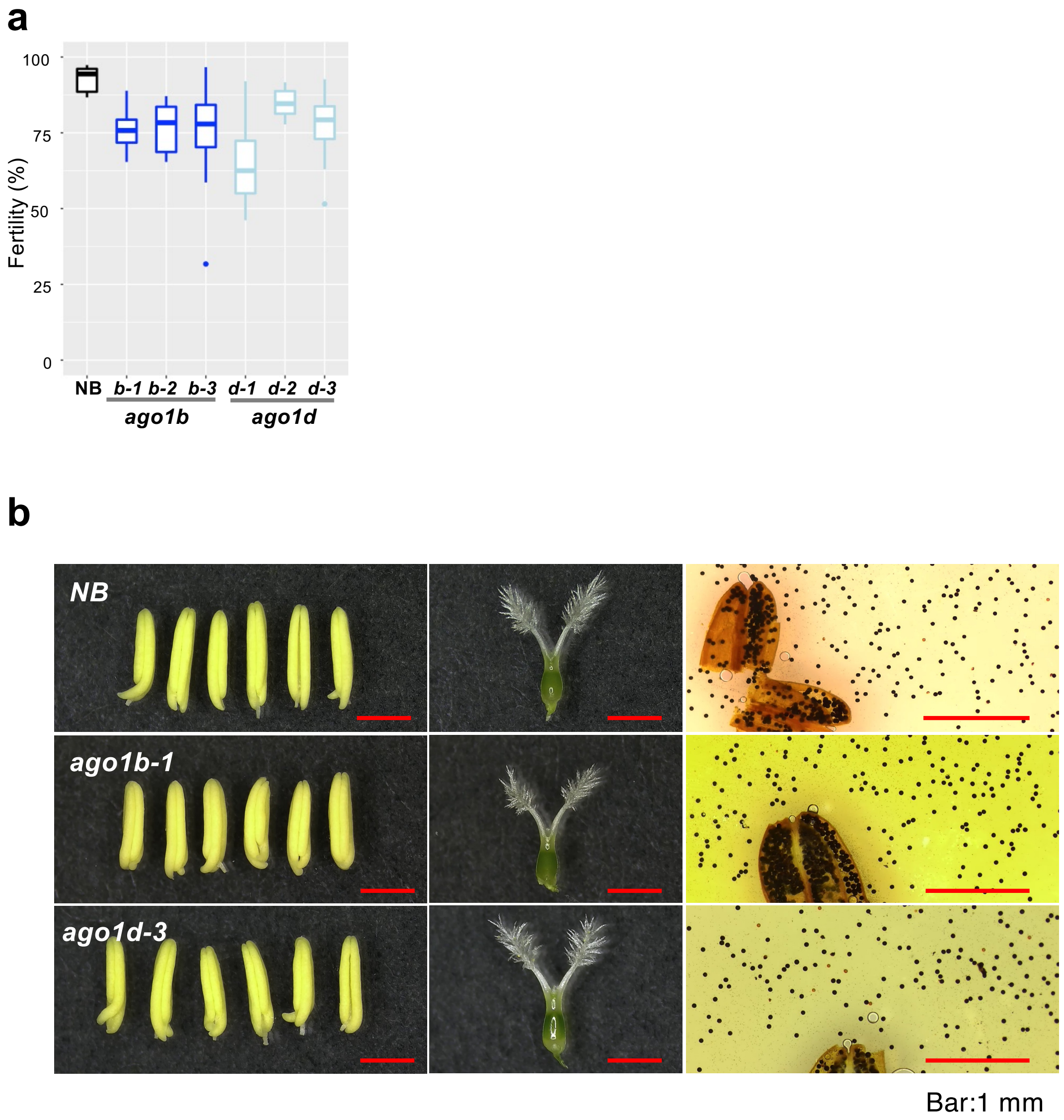


**Supplementary Figure 2. a**. Fertility of Nipponbare (NB), *ago1b-1* (*b-1*), *ago1b-2* (*b-2*) *ago1b-3* (*b-3*), *ago1d-1* (*d-1*), *ago1d-2* (*d-2*), and *ago1d-3* (*d-3*). Data shown in box plots are from more than three biological replicates. Student's *t* test. **b**. Mature anthers (left), pistil (middle), and pollen grains (right) of Nipponbare (control), *ago1b-1*, and *ago1d-3*. The single mutants, *ago1b-1* and *ago1d-3*, showed normal anthers and pistils compared to the those of Nipponbare. Furthermore, mature pollen grains of *ago1b-1* and *ago1d-3* were also stained with iodine-potassium iodide.

Supplementary Figure 3

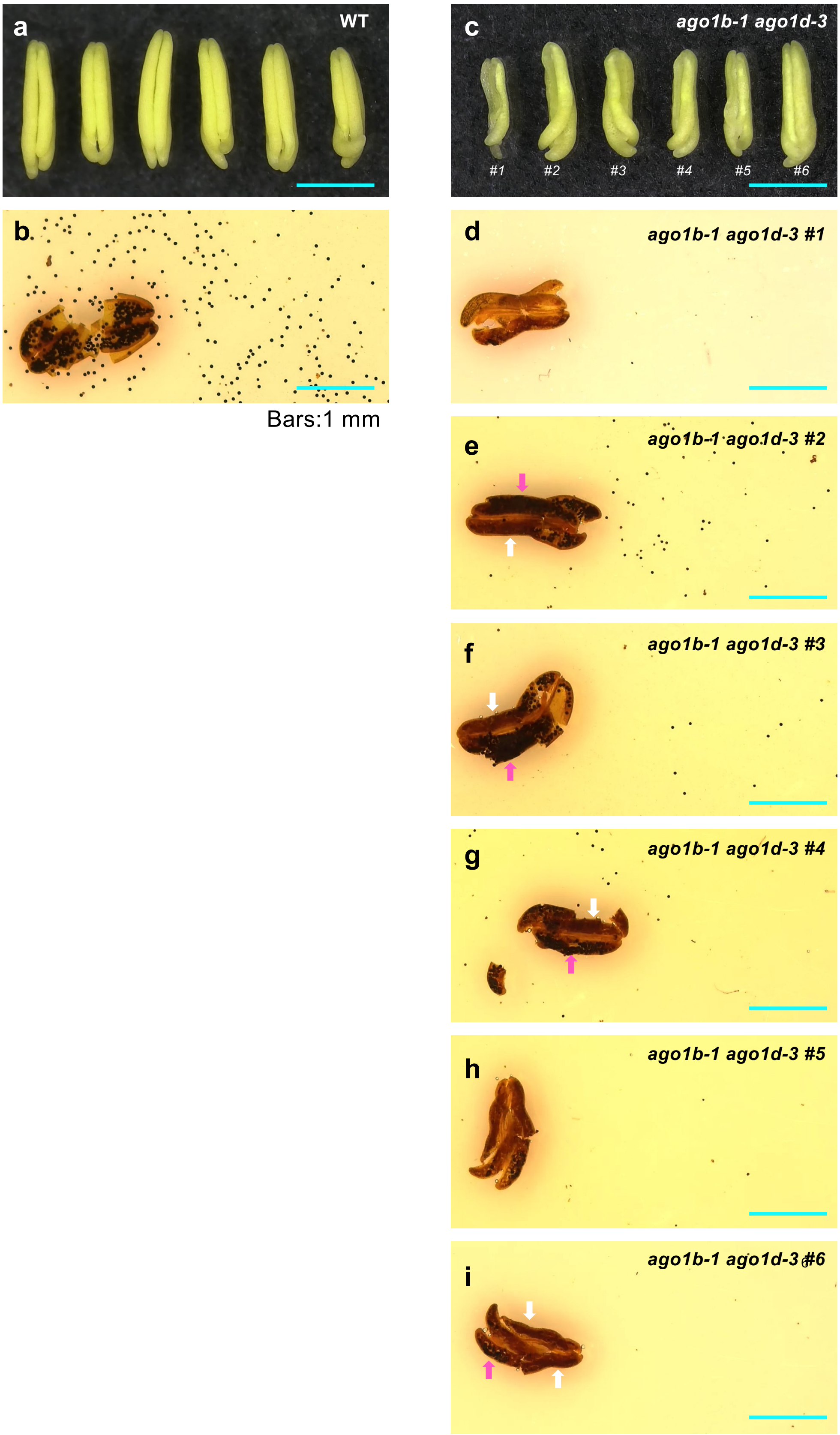


**Supplementary Figure 3. a**. Anthers of WT. **b**. Starch staining of pollen from WT anthers. **c**. Anthers of *ago1b-1 ago1d-3* double mutants with several abnormal shapes from severe to mild types (#1–6). **d–i**. Starch staining of pollen from a severely abnormal and semi-abnormal anthers of the *ago1b-1 ago1d-3* double mutant (#1–6). Non-staining pollens reflect the abnormality of pollen activity and development (**d and h**). Semi-abnormal anthers of the double mutant contain non-staining pollens (white arrows) as well as stained pollen grains (magenta arrows) (**e, f, g, and i**). Most of the stained pollen grains were trapped in the anthers of the *ago1b-1 ago1d-3* double mutant (**e, f, g, and i**), perhaps due to defects of somatic anther wall developments. Scale bars are 1 mm.
